# Myosin 1b Flattens and Prunes Branched Actin Filaments

**DOI:** 10.1101/2020.02.26.966663

**Authors:** Julien Pernier, Antoine Morchain, Valentina Caorsi, Aurélie Bertin, Hugo Bousquet, Patricia Bassereau, Evelyne Coudrier

## Abstract

Motile and morphological cellular processes require a spatially and temporally coordinated branched actin network that is controlled by the activity of various regulatory proteins including the Arp2/3 complex, profilin, cofilin and tropomyosin. We have previously reported that myosin 1b regulates the density of the actin network in the growth cone. Using *in vitro* F-actin gliding assays and total internal reflection fluorescence (TIRF) microscopy we show in this report that this molecular motor flattens the Arp2/3-dependent actin branches up to breaking them and reduces the probability to form new branches. This experiment reveals that myosin 1b can produce force sufficient enough to break up the Arp2/3-mediated actin junction. Together with the former *in vivo* studies, this work emphasizes the essential role played by myosins in the architecture and in the dynamics of actin networks in different cellular regions.

**Short summary:** Using *in vitro* F-actin gliding assays and total internal reflection fluorescence (TIRF) microscopy we show that myosin flattens the Arp2/3-dependent actin branches up to breaking them and reduces the probability to form new branches

## Introduction

Many motile and morphological processes involve spatially and temporally coordinated branched actin-filament (F-actin) networks. Polymerization and assembly of branched networks are controlled by the activity of regulatory proteins including the Arp2/3 complex (Blanchoin et al., 2014), profilin (Pernier et al., 2016), cofilin (Chan et al., 2009) and tropomyosin (Blanchoin et al., 2001). In addition, several myosin motors have been reported to control actin architecture, either bound to cell membranes such as myosins 1 (see below), 6 (Loubéry et al., 2012; Reymann et al., 2012; Ripoll et al., 2018) and 10 (Nagy et al., 2008) or associated to F-actin only, such as myosin II (Haviv et al., 2008; Mendes Pinto et al., 2012; Reymann et al., 2012).

Myosins1 are single-headed motors containing three domains: a N-terminal motor domain that coordinates ATP hydrolysis, actin binding and force generation, a light chain binding domain (LCBD) that binds calmodulin, and a tail region with a highly basic C-terminal tail homology 1 (TH1) (McIntosh and Ostap, 2016) (Fig.1A). In contrast to non-muscle myosinII, a pleckstrin homology (PH) motif in the TH1 domain targets myosins1 to phosphoinositides in membranes (McConnell and Tyska, 2010) (Fig.1A). In *S cerevisiae,* myosin1 (Myo5) induces actin polymerisation depending on the Arp2/3 complex (Sirotkin et al., 2005). This myosin1 exhibits a SH3 domain at its C-terminus that binds the WASP yeast homologue followed by an acidic motif binding Arp2/3 (Evangelista et al., 2000). Studies in Xenopus oocytes suggest that myosin1c and myosin1e control cortical granule exocytosis and the subsequent compensatory endocytosis by directly regulating actin polymerization (Schietroma et al., 2007; Sokac et al., 2006; Yu and Bement, 2007). Several members of this myosin family in mammals are also involved in actin networks’ organisation. Myosin1c stabilizes actin around the Golgi complex (Capmany et al., 2019). Myosin1b (Myo1b) regulates neuronal polarization and axon elongation by limiting the extension of branched actin networks and favoring actin bundles in growth cones (Iuliano et al., 2018). However, the mammalian myosins1 lack a SH3 domain or an acidic motif that could interact with Arp2/3. We have recently reported that single F-actins depolymerize when they slide on immobilized Myo1b (Pernier et al., 2019) but how mammalian myosins1 reorganize branched actin network architecture remains to be explored.

**Figure 1:**
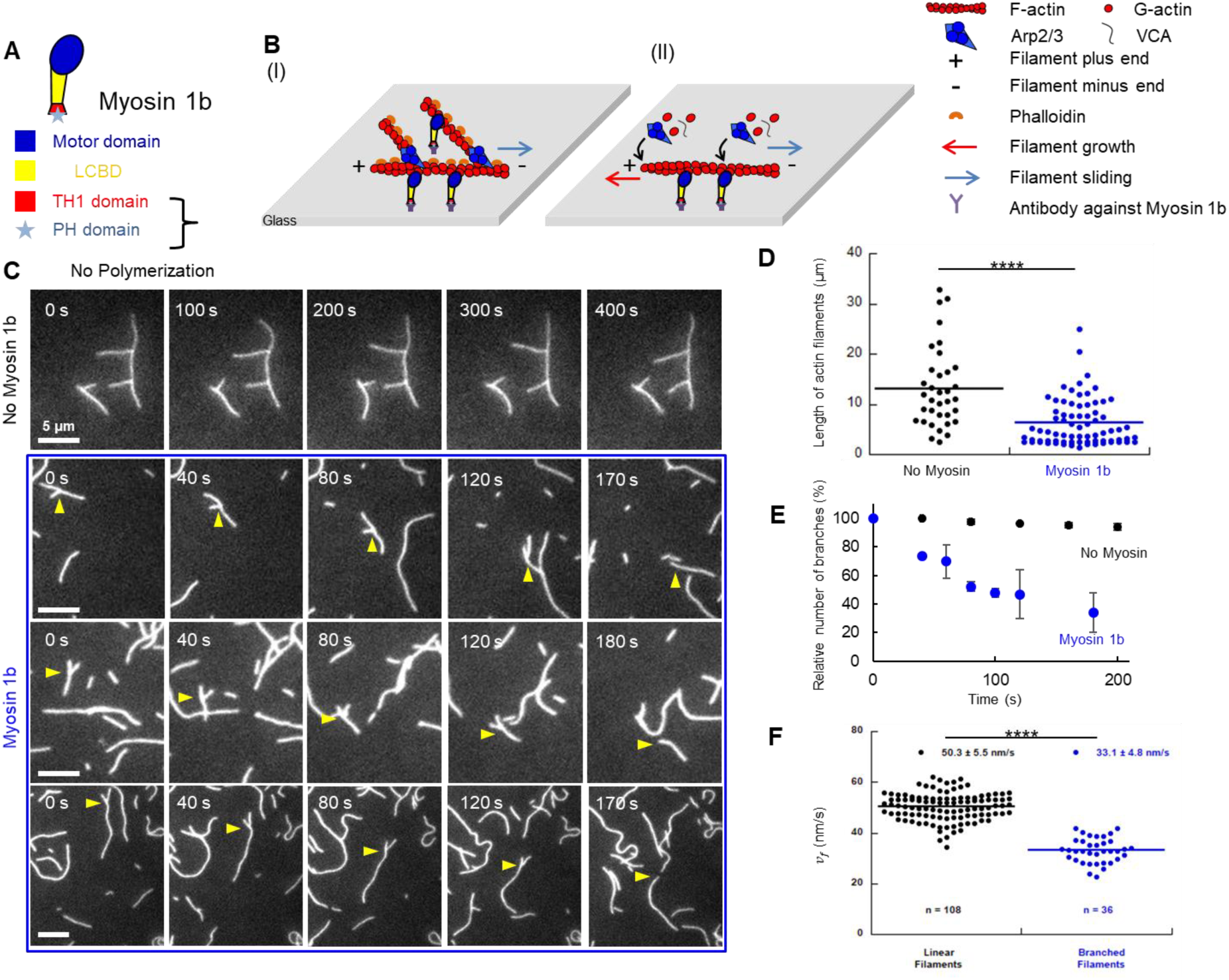
Sliding on Myo1b reduces stabilized F-actin branching. **(A)** Myo1b domain organization. **(B)** Sketch illustrating the gliding assays of phalloidin-stabilized branched (I) and polymerizing and branching (II) F-actin sliding on Myo1b anchored to coverslip. **(C)** Representative time-lapse images of phalloidin-stabilized branched F-actin sliding or not on Myo1b (MovieS1). Yellow arrows point to a single branch. Scale bar, 5μm. (**D)** Distribution of F-actin (mother and daughter) lengths at t=160s, in one field corresponding to MovieS1 (no myosin and Myo1b, 2^nd^ from left). A two-tailed t test (p=2.08×10^-5^) shows a significant difference. (**E)** Mean relative number of branches along F-actin (Methods) normalized by N0, the branch number at t=0, over time in the absence (N0=60, 2 movies) or sliding on Myo1b (N_0_=38, 2 movies). s.e.m. are represented. (**F)** Dot plot of sliding velocities *v_f_* of stabilized F-actin in the presence of Arp2/3 showing or not branches, analyzed on the same movies (5 movies). Number of analyzed filaments and mean-values ± s.e.m. are indicated. A two-tailed t-test (p=2.89×10^-27^) shows a significant difference between data sets. ****p < 0.0001.

Here, we used *in vitro* F-actin gliding assays and total internal reflection fluorescence (TIRF) microscopy to study how the organization (I) and the dynamics (II) of stabilized (I) and polymerizing (II) branched F-actin are modified when sliding on full-length Myo1b immobilized on a substrate (Fig.1B). We compared the effect of Myo1b in these experimental conditions with another myosin, the muscle myosin II (Myo2) that moves linear F-actin five-fold faster (Pernier et al., 2019), keeping in mind that these experimental conditions mimic the physiological topology reported for Myo1b but not for Myo2.

## Results and Discussion

### Branch density of stabilized F-actin decreases when sliding on Myo1b

We first polymerized F-actin in solution, in the presence of the constitutively active C-terminal domain of N-WASP (VCA) and Arp2/3 complex (Methods) (Pernier et al., 2016), to generate branched filaments and stabilized them with phalloidin. We analyzed the filaments’ architecture and movement when sliding on Myo1b bound to the glass substrate at a density of a 8000/μm^2^ (Pernier et al., 2019) (Fig.1C, movieS1). Without Myo1b and with the same surface treatment, the branches are stable at least up to 10 minutes (MovieS1, Figs.1C, 1E). In contrast, when filaments slide on motors, we observe large variations of the orientation of the branches relative to the mother filaments, already visible 1 or 2 minutes after gliding (eg MovieS1, Fig.1C and Fig.2C) and detachment of branches from the mother at the level of the branch attachment (MovieS1, Fig.1C, last frames). Quantification of F-actin length distribution after 160 sec shows an important and significant (p=2.08×10^-5^) increase of short filaments when gliding on Myo1b (Fig.1D), while the total length (including mother, daughter and single filaments) is conserved from the initial frame to the last one (457 and 459 μm respectively, (N=34 at t = 0sec, N=35 at t= 160sec), similarly to the control (448 and 459 μm). In average, the filament length is 13.1μm without Myo1b and 6.2μm in its presence (Fig.1D). Since filaments are stabilized with phalloidin, reduction of their length when sliding on Myo1b cannot be due to its actin depolymerase activity (Pernier et al., 2019). It is not due to a severing activity of Myo1b either: we have observed that 100% of the breaks occur at the base of the shorter (daughter) filament within TIRF resolution (1 pixel=160nm). In contrast, severing would occur everywhere along mother and daughter filaments. Moreover, we did not detect severing on stabilized linear filaments (Pernier et al., 2019). Thus, our observations rather suggest that sliding on Myo1b induces branches’ detachment.

**Figure 2:**
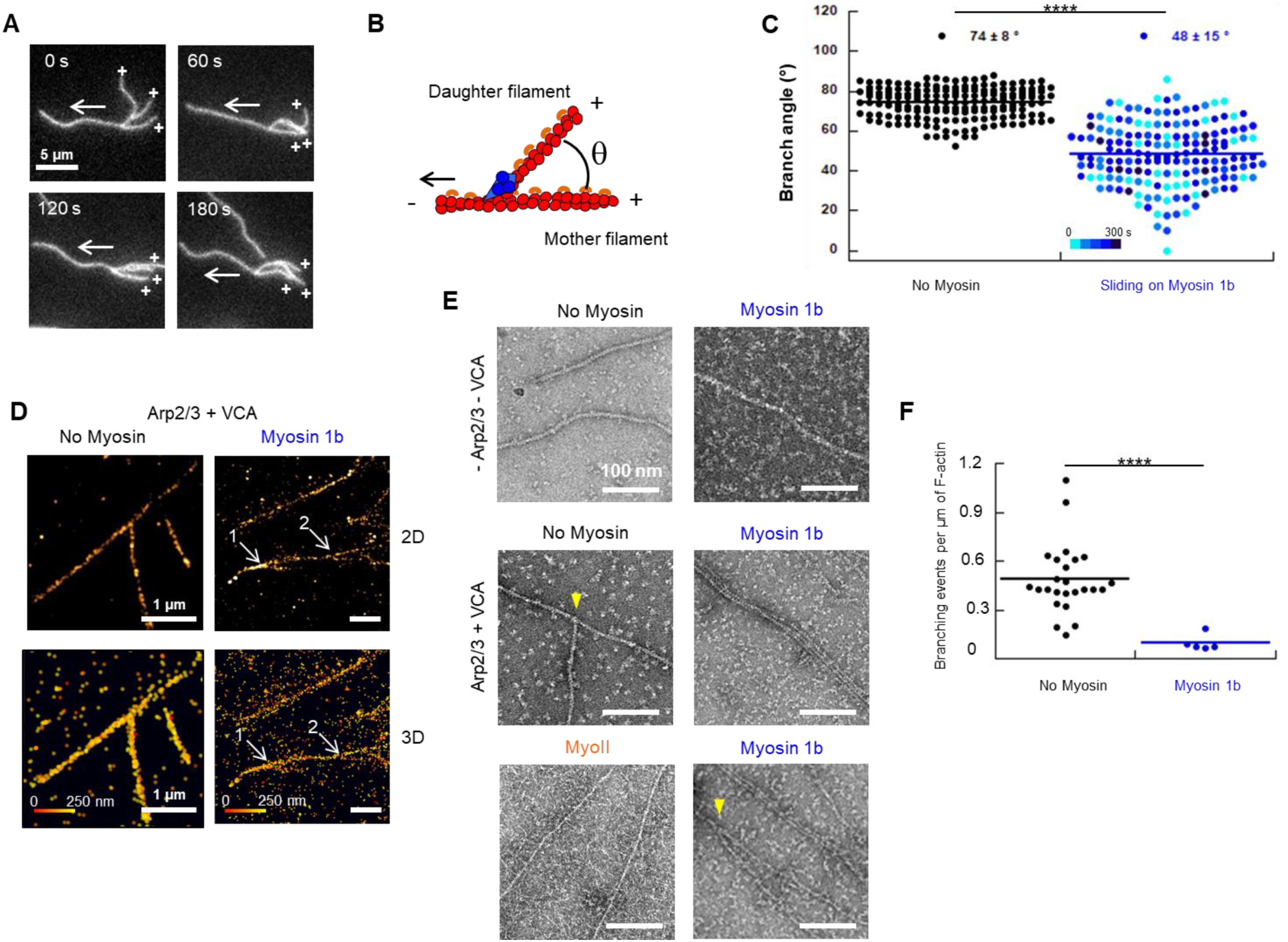
Myo1b reduces stabilized F-actin angles in branched structures. **(A)** Time-lapse images of stabilized branched F-actin, sliding on Myo1b, showing a change of the angle between mother and daughter filaments (MovieS3). Crosses indicate barbed ends and white arrows the sliding direction. Scale bar, 5μm. **(B)** Representation of the θ angle between mother and daughter filaments. **(C)** Dot plot of θ angle without myosin (12 filaments, 168 images) or sliding on Myo1b (19 filaments, 168 images). Blue scale indicates the time since acquisition started. A two-tailed t-test (p=6.16×10^-51^) shows a significant difference. **(D)** STORM images of stabilized branched F-actin in 2D (**top**) or 3D (**bottom**) without or with Myo1b. They correspond to the squares in Fig.S1A. Color code indicates the height (z). Without Myo1b, branch and mother filaments are in the same plane and attached. On Myo1b, branches #1 and #2 exhibit a θ angle much lower than 70°, but are physically connected to it. Note that branch #2 is in a lower plane. **(E)** Electron microcopy images of stabilized F-actin branched or not, recorded after gliding 10 min or not on Myo1b or Myo2 and after negative staining. These images correspond to the squares in Fig.S1B. Scale bars, 100nm. **(F)** Dot plot representing the number of branches per μm of F-actin, without myosin (N=25) or sliding on Myo1b (N=5), quantified from electron microscopy images. A two-tailed t-test (p=9.78×10^-9^) shows a significant difference. ****p < 0.0001.

As a comparison, we used Myo2 that leads to a five-fold faster linear F-actin sliding (Pernier et al., 2019). At the same surface density, no branches on filaments gliding on Myo2 but F-actin fragments were observed, likely due to a fast detachment occurring before image acquisition started, i.e. one minute after injection of branched filaments in the chamber (MovieS2). Since we aim at characterizing the debranching process by myosins, we focus on Myo1b in the following, given the time scale of our experiments.

We next quantified the relative variation of the number of branches along F-actin over time. The branch number decreases by five-fold within 3 minutes on Myo1b (Fig.1E). The presence of branches reduces the gliding velocity, from 50.3±5.5nm/sec for filaments without branches as we previously reported (Pernier et al., 2019; Yamada et al., 2014) down to 33.1±4.8 nm/s for branched F-actin (Fig.1F).

### Sliding on Myo1b flattens the branches on the actin filaments

Since we observed variations of the branches’ orientation (Figs.1C, 2A, MoviesS1, S3), we measured the θ angle between mother and daughter filaments (Fig.2B) on a large collection of snapshots, without or with Myo1b. In average, θ=74±8° in control conditions, in agreement with previous reports (Mullins et al., 1998; Svitkina and Borisy, 1999) (Fig.2C). In contrast, with Myo1b, θ decreases to 48±15°, with a large spreading of values down to almost zero (Fig.2C). This large spreading is maintained over 5 minutes’ observation (Fig.2C, blue scale), reflecting a large flexibility of the branched filaments architecture when sliding on Myo1b. In many instances the branch flattens on the mother (MoviesS1, S3) and the angle decays to zero; the branch falls back towards the main filament axis in the direction opposite to the movement. Using 3D STORM, we confirmed that even when θ is close to zero, the branch remains attached to the mother filament, and is not juxtaposed to it (Fig.2D, Fig.S1A).

We next analyzed the architecture of the actin networks with electron microscopy (Fig.2E, Fig.S1B). Figure 2E shows a representative image of a branched filament without Myo1b with θ=67°, of a bundle of 3 parallel filaments and of a branch with a reduced angle with Myo1b. In average, we measured 0.48 branches/μm without Myo1b (N=25 fields of 2.5X1.7μm^2^), and 0.1 branches/μm with Myo1b (N=5) (Fig.2F). In agreement with our time-lapse observations (MovieS2), we observed neither branches nor bundles with Myo2 (Fig.2E).

All together, these observations highlight Myo1b capability to modify the architecture of branched stabilized actin networks, to reduce the branching angle down to zero, in the direction opposite to the movement and to even break Arp2/3-mediated branches.

### Sliding on Myo1b reduces branching dynamics and induces debranching

We next investigated the effect of Myo1b on dynamical filaments that polymerize and branch, using bulk pyrene-actin fluorescence or TIRF microscopy assays. As previously reported for linear F-actin, Myo1b in solution does not change polymerization of branched F-actin (Fig.3A) (Pernier et al., 2019). Using TIRF assay, we observed the growth of long and straight branches along the growing mother filament in the absence of myosin, as expected (Fig.3B, MovieS4). In contrast, when sliding on Myo1b, the growing filaments appear very distorted. As previously reported in the absence of Arp2/3 (Pernier et al., 2019), Myo1b also reduces the actin polymerization rate in these conditions. It decreases from 7.37±0.57 su/s (N= 60) without myosin (Pernier et al., 2019) down to 5.32 ±0.66 su/s (N= 25) with Myo1b. Branches also form over time when sliding on Myo1b but they frequently detach and run away (Fig.3B, MovieS4, Fig. S2A,). We normalized the branch number measured at different times by the corresponding F-actin total length (Fig.S2C). The number of branches formed per μm is time-independent but is reduced by a factor 2 (0.11±0.02.μm^-1^) when sliding on Myo1b as compared to the control (0.24±0.02.μm^-1^) (Fig.3C). The cumulative number of debranching events increases over time when sliding on Myo1b while none occurs in control conditions (Fig.S3B). When normalized by the total F-actin length (Fig.3D), the density of debranching events is 0.05±0.01.μm^-1^. The observed normalized branch number (Fig.3C) corresponds to the difference between the normalized branch numbers effectively formed (Fig. S2D) and those that detach (Fig.3D). Measurements in Fig.3C and Fig.3D show that the effective number of newly formed branches is equal to 0.16±0.02.μm^-1^, lower than in the control (Fig.S2D). This indicates that sliding of F-actin on Myo1b, besides reducing polymerization rate, induces debranching and also reduces the probability of generating new branches.

**Figure 3:**
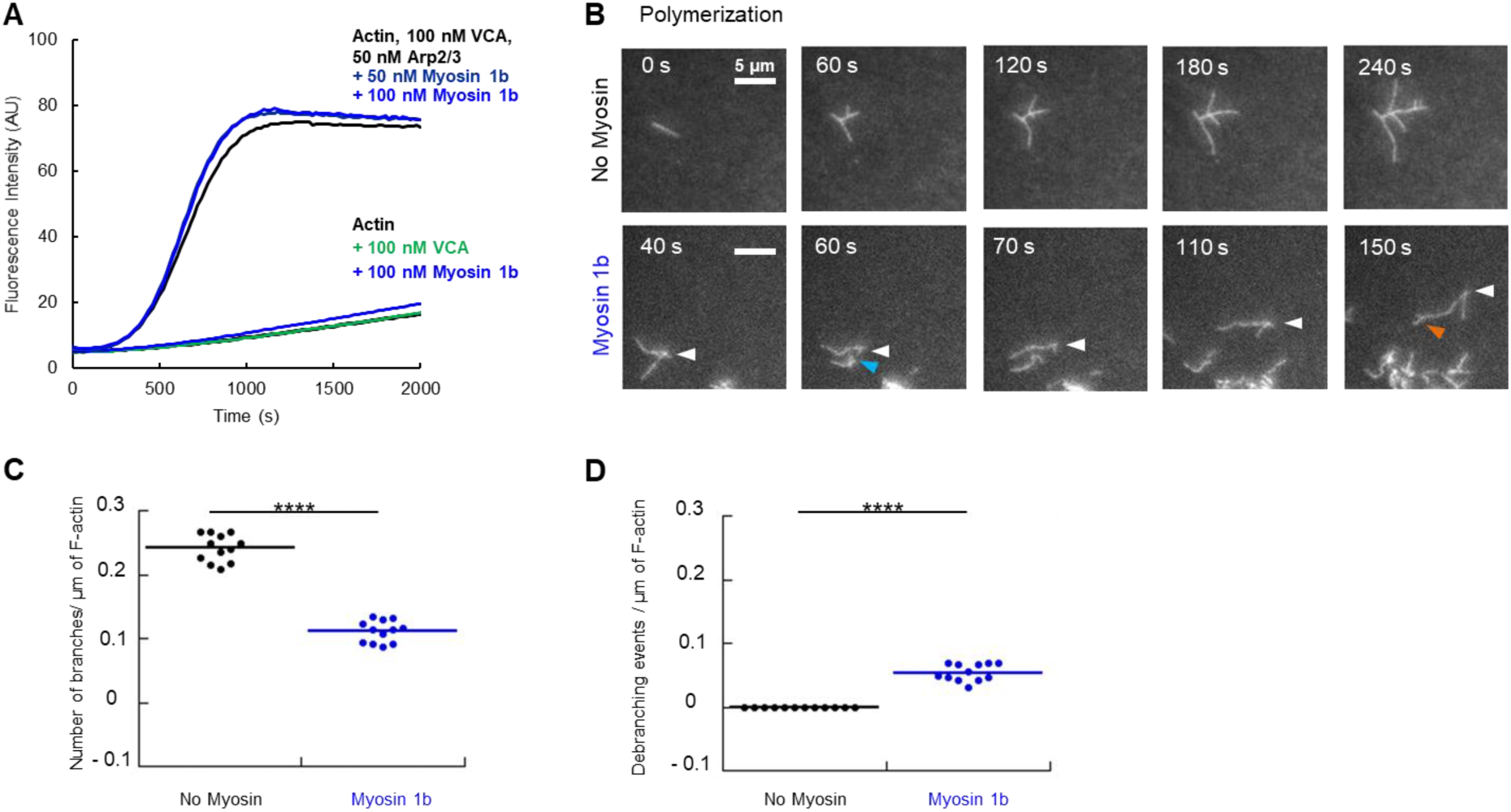
Sliding on Myo1b decreases branching of polymerizing actin filaments. **(A)** Effect of Myo1b on branched F-actin polymerization using pyrene assay. **(B)** Time-lapse images of polymerizing and branching F-actin without or sliding on Myo1b at 2mM ATP (MovieS4). Scale bar, 5μm. White, Cyan and orange arrows point to a mother filament debranching and branching events respectively. **(C)** Numbers of branches per μm of F-actin, without myosin or sliding on Myo1b. They are obtained from the total number of branches detected at different times (Fig.S2A) normalized by the corresponding total F-actin length (Fig.S2C) (3 movies). A two-tailed t-test (p=1.47×10^-13^) shows a significant difference. **(D)** Numbers of debranching events per μm of F-actin, without or with Myo1b, obtained from the same normalization as in (C), using (Fig.S2B) and (Fig.S2C). A two-tailed t-test (p=2.61×10^-8^) shows a significant difference.

It is generally accepted that both actin network types, branched and parallel depending on Arp2/3 complex and formin respectively, compete for the same G-actin pool (Chesarone and Goode, 2009). More recently it was shown that accessory proteins such as profilin or capping protein can interfere and favor the polymerization of one or the other network (Rotty et al., 2015; Suarez et al., 2015). In this report we show that beside the accessory proteins, Myo1b and Myo2 also remodel the architecture of branched actin networks when attached to a substrate. This is not the case when myosins are free in solution and cannot exert forces. Bieling and colleagues (Bieling et al., 2016) suggested that compressive forces on a branched actin gel alter the internal architecture of the network, increasing its density. Recent work using microfluidics (Pandit et al., 2020) confirms that Arp2/3-dependent branches can break when forces in the pN range are applied. In our experiments on stabilized and dynamical F-actin, due to the geometry of the Arp2/3-generated branch, the mother filament and the branches are propelled by myosins in divergent directions, which induces friction forces on the filaments (Fig.4). This explains the reduction of sliding velocity in the presence of branches. Moreover, combination of the friction force (*F_fric_*) on the branch due to mother filament sliding and of the force produced by motors on this branch (*F_mot_*) results on a force **F** at the filament extremity, and thus on a torque that reduces the θ-angle (Fig.4). Consequently, the Arp2/3 junction is submitted to a stress and depending on the stiffness of the Arp2/3/F-actin junction, the branch might break either at the junction or next to it.

**Figure 4:**
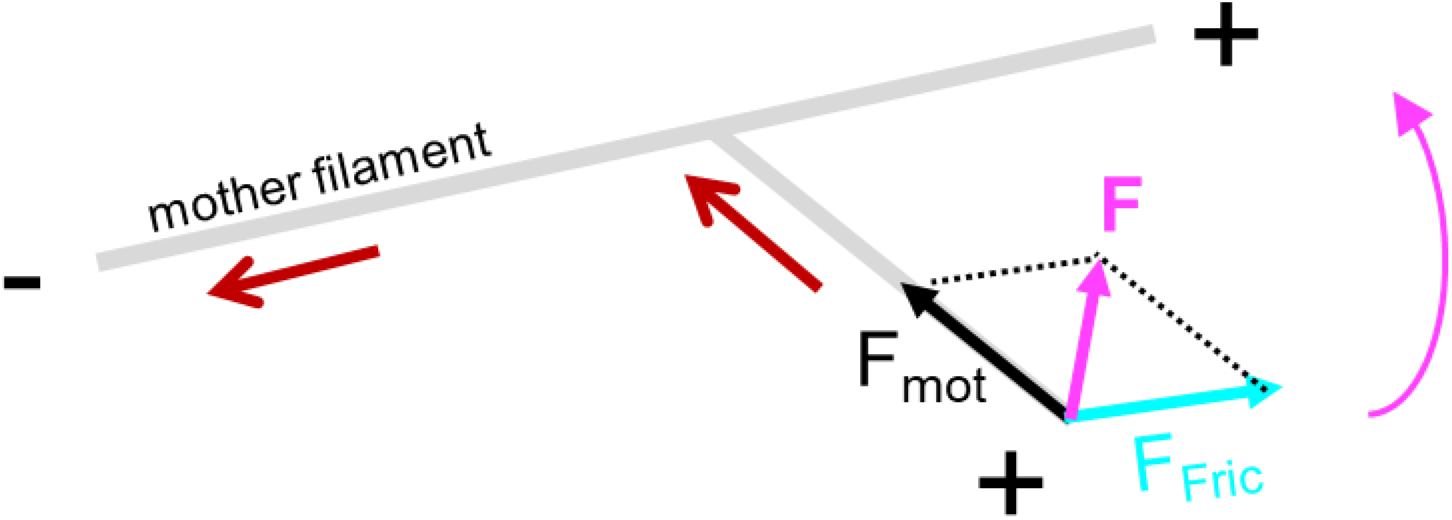
Scheme of the forces exerted on a daughter filament. Mother filament sliding (red arrow) induces a friction force (*F_fric_*) on the daughter filament. *F_fric_* combined to the force generated by the motors on the branch (*F_mot_*) results in a total force ***F*** at the extremity of the filament and thus in a torque leading to the reduction of the angle between both filaments (magenta arrow).

We have also shown that the probability of forming new branches on the side of a mother filament is reduced when sliding on myosins. Tropomyosin (Blanchoin et al., 2001), cofilin (Chan et al., 2009) and coronin (Cai et al., 2008) inhibit the polymerization of Arp2/3-dependent actin branches by competing with the Arp2/3 complex for their binding site along F-actin. Similarly, Myo1b that also binds along F-actin (Mentes et al., 2018) might compete with Arp2/3 for the same binding site.

In cells, Myo1b is bound to membranes. The Ostap group and us (Pernier et al., 2019; Pyrpassopoulos et al., 2012) have reported that even when Myo1b is bound to fluid lipid bilayers, it can propel F-actin due to the high membrane viscosity (Pernier et al., 2019). Thus, it is very likely that Myo1b also debranches and reorganizes the architecture of branched actin networks at the plasma membrane. This is supported by our *in vivo* observations that Myo1b is required for the formation of filopodia induced by EphB2 receptor stimulation (Prospéri et al., 2015) and controls the extension of branched actin networks in growth cones (Iuliano et al., 2018). By debranching actin networks nearby the plasma membrane, Myo1b may favor the formation of linear and parallel filaments in growth cones and thus facilitate filopodia initiation upon EphB2 receptor stimulation. Eventually, we showed that Myo1b can pull membrane tubules along actin bundles (Yamada et al., 2014) reminiscent of the Myo1b-dependent tubule elongation at the trans-Golgi network (Almeida et al., 2011). An additional role for Myo1b in this cellular region could be to debranch the Arp2/3 network to form actin structures required for it to pull tubules. Other Myosins1 were reported to influence actin organization *in vivo* (Capmany et al., 2019; Schietroma et al., 2007; Sokac et al., 2006; Yu and Bement, 2007), however their motor properties and mechanosensitivity differ from Myo1b (McIntosh and Ostap, 2016), thus they might act differently on actin branches. Myo2 also inhibits branched actin structures. It is important to stress that this motor is not membrane-bound but associated with contractile fibers. Whether or not Myo2 *in vivo* contributes to formation of contractile fibers by debranching Arp2/3 actin network remains to be studied.

This work, together with former *in vivo* studies, reveals the essential role played by Myo1b in the architecture and in the dynamics of actin networks at the plasma membrane and on cellular compartments. Considering its motor activity that produces F-actin motion, we can envisage that Myo1b is also involved in general actin movements and fluxes at larger scales.

## Acknowledgments

We thank B. Goud and J.F. Joanny (Institut Curie) for insightful discussions, C. Le Clainche (I2BC, Gif-sur-Yvette, France) for providing actin and for critically reading the manuscript, A-S. Mace for developing a macro to analyze the movies, G. Romet-Lemonne and L. Blanchoin for carefully reading the manuscript. The authors greatly acknowledge the Cell and Tissue Imaging (PICT-IBiSA), Institut Curie, member of the French National Research Infrastructure France-BioImaging (ANR10-INBS-04). This work was supported by Institut Curie, Centre National de la Recherche Scientifique, the European Research Council project 339847. J.P. was funded by the ERC project 339847. P.B. and E.C.’s groups belong to the CNRS consortium CellTiss, the Labex CelTisPhyBio (ANR-11-LABX0038) and Paris Sciences et Lettres (ANR-10-IDEX-0001-02).

## Author contributions

J.P., P.B and E.C. designed the study. J.P. and A.M. performed TIRF experiments and analyzed data; V.C. performed the STORM experiments and analyzed data; A.B. performed the EM experiments; H.B. purified myosin 1b; J.P, P.B and E.C., wrote the paper.

## Declaration of interests

The authors declare no competing interests.

## Supplemental Information

**Figure S1:**
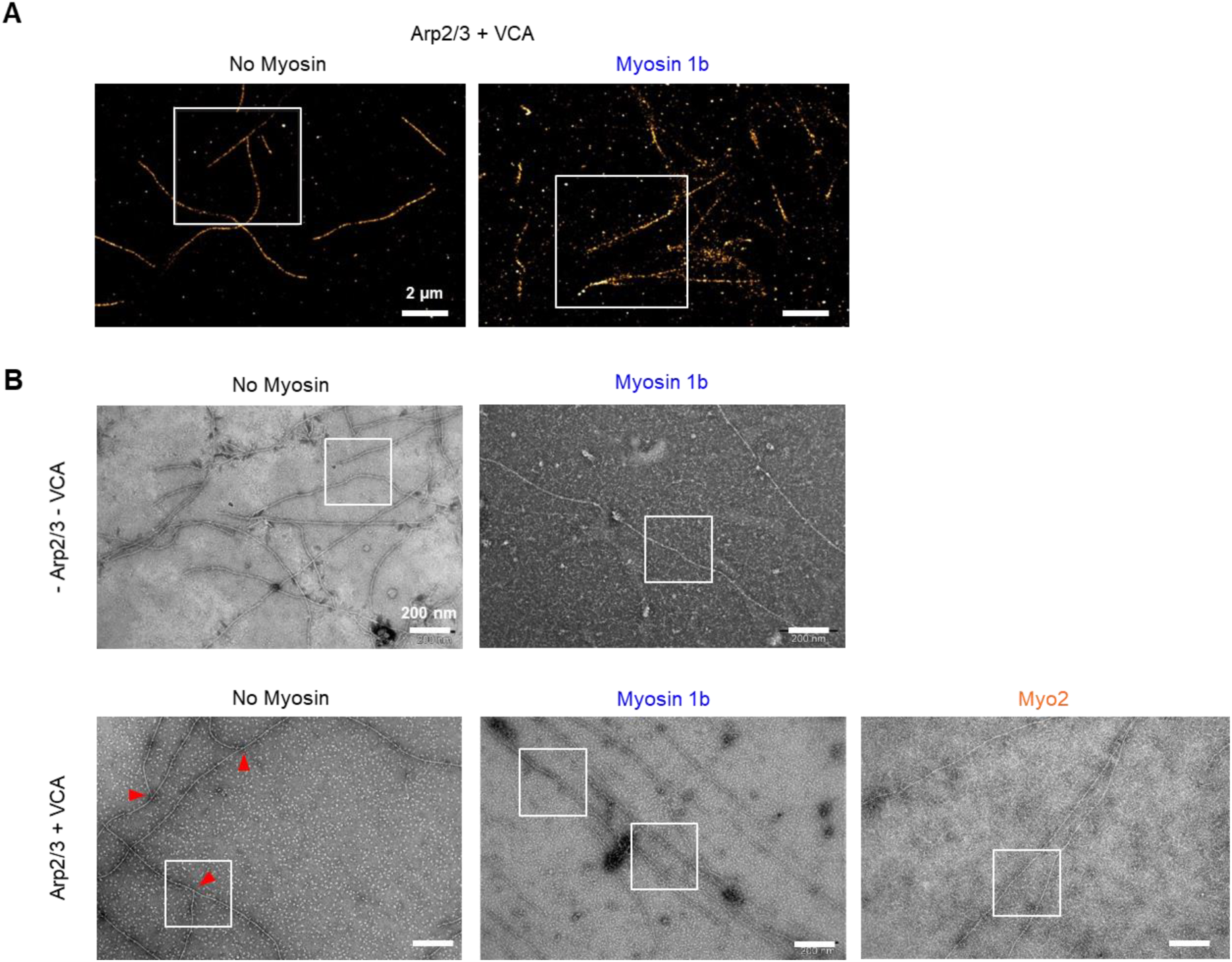
Large fields corresponding to the zooms (white frames) shown: (A) in Fig. 3D with STORM and (B) in Fig. 3E with electron microscopy. In (B), red triangles indicate branched junctions. Scale bars 2 μm (A) and 200 nm (B).

**Figure S2:**
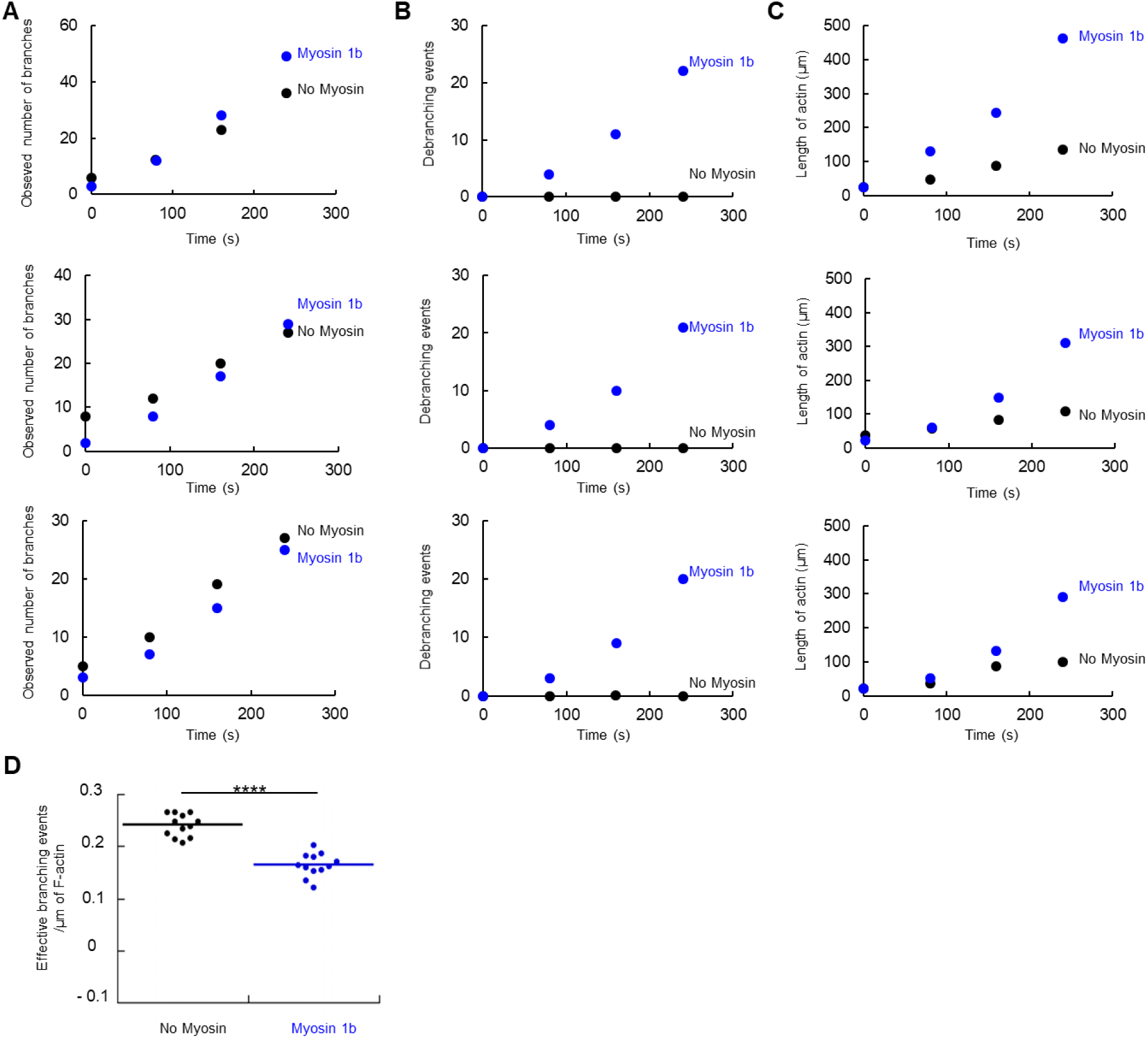
Data corresponding to Fig. 3C and 3D. **(A)** Observed number of branches, **(B)** total debranching events, **(C)** total length of actin, over time and on three different fields of 80×80 μm^2^. **(D)** Effective branching events per μm of F-actin, calculated from the sum of the observed branches number (Fig. 3C) and debranching events (3D), without or with Myo1b. A two-tailed t-test (p=1.51×10^-8^) shows a significant difference. ****p < 0.0001.

**Movie S1: Stabilized branched actin filaments sliding along glass-anchored myosin 1b (corresponding to Fig. 2).**

Branched filaments stabilized with phalloidin-Alexa547 in the absence of myosin (left) or sliding on Myo1b at high density (8000 μm ^-2^). Scale bar 5 μm. Time in s. Wide field movies corresponding to time-lapse images shown in Fig. 1C.

**Movie S2: Stabilized branched actin filaments sliding along Myo2.**

Branched filaments stabilized with phalloidin-Alexa547 sliding on Myo2. The movie starts 1 min after injection of branched filaments in the chamber. Scale bar, 5 μm. Time in s.

**Movie S3: Effect of myosin 1b on organization of branched structures (corresponding to Fig. 2A).**

Branched filaments stabilized with phalloidin-Alexa547 sliding on Myo1b. Scale bar, 5 μm. Time in s.

**Movie S4: Polymerizing and branching actin filaments sliding along glass-anchored Myo1b (corresponding to Fig. 3).**

Polymerized and branched actin filaments in the absence of myosin or sliding on Myo1b at high density (8000 μm ^-2^). Scale bar, 5 μm. Time in s. Wide field movies corresponding to time-lapse images shown in Fig. 3B. White arrow points to a mother filament. The appearance over time of new actin filaments on the right side of the movie likely results from the high affinity of these filaments for the surface coated with Myo1b.

## STAR Methods

### Protein purification

Actin was purified from rabbit muscle and isolated in monomeric form in G buffer (5 mM Tris-HCl, pH 7.8, 0.1 mM CaCl2, 0.2 mM ATP, 1 mM DTT and 0.01% NaN3). Actin was labeled with Alexa 594 succimidyl ester-NHS (Ciobanasu et al., 2014).

Myosin II was purified from rabbit muscle as previously described (Pollard, 1982).

Arp2/3complex was purified from bovine brain (Egile et al., 1999).

Expression and purification of Myosin 1b: FLAG-myo1b was expressed in HEK293-Flp-In cells regularly tested for contamination, cultured in Dulbecco’s modified Eagle medium supplemented with 10% fetal bovine serum and 0.18 mg ml^-1^ hygromycine in a spinner flask at 37 °C under 5% CO_2_, and collected by centrifugation (1,000 g, 10min, 4 °C) to obtain a 4–5 g of cell pellet as described by Yamada et al. (Yamada et al., 2014). The pellet was lysed in FLAG Trap binding buffer (30 mM HEPES, pH 7.5, 100 mM KCl, 1 mM MgCl_2_, 1mM EGTA, 1 mM ATP, 1 mM DTT, 0.1% protease inhibitor cocktail (PIC), 1% Triton X-100) for 30 min at 4 °C and centrifuged at 3,400 g for 10 min at 4 °C. The collected supernatant was then ultracentrifuged (250,000 g, 60 min, 4 °C). The solution between pellet and floating lipid layer was incubated with 150 μl of anti-FLAG beads for 2h at 4 °C. The beads were collected by centrifugation (1,000 g, 5 min, 4 °C). After a washing step, FLAG-myo1b was then eluted by incubating with 0.24 mg ml^-1^ of 3X FLAG peptide in 300 μl elution buffer (binding buffer without Triton X-100 supplemented with 0.1% methylcellulose) for 3 h at 4 °C. After removal of the beads by centrifugation (1,000 g, 3 min, 4 °C), the protein solution was dialyzed against elution buffer overnight at 4 °C to remove the 3X FLAG peptide. Myo1b was fluorescently labeled using Alexa Fluor 488 5-SDP ester (Yamada et al., 2014). Inactivated Myo1b was removed by ultracentrifugation (90,000 rpm, 20 min, 4 °C) with 10 μM F-actin in presence of 2 mM ATP. Inactivated Myo1b was then dissociated from F-actin by incubating the pellet collected after untracentrifugation in elution buffer (30 mM HEPES, pH 7.5, 100 mM KCl, 1 mM MgCl_2_, 1mM EGTA, 1 mM ATP, 1 mM DTT and 0.1% methylcellulose) supplemented with 1 M NaCl and collected in the supernatant after a second centrifugation (90,000 rpm, 20 min, 4 °C).

### Actin polymerization assays using pyrenyl assay

Actin polymerization kinetic experiments were based on fluorescence change of pyrenyl-labeled G-actin (λ_exc_=365 nm, λ_em_=407 nm). Experiments were carried out on a Safas Xenius spectrofluorimeter (Safas, Monaco). Polymerization assays were performed in F-buffer (5 mM Tris-HCl, pH 7.8, 100 mM KCl, 1 mM MgCl_2_, 0.2 mM EGTA, 0.2 mM ATP, 1 mM DTT, 0.01% NaN_3_) in the presence of 100 nM VCA, 50 nM Arp2/3 and 2 μM Actin 10% pyrenyl-labeled).

### TIRF microscopy assays and data analysis

The kinetics of single filament assembly Coverslips and glass slides were sequentially cleaned by sonication with H2O, ethanol, acetone for 10 min, then 1M KOH for 20 min and H2O for 10 min. Flow chambers were assembled with a coverslip bound to a glass slide with two parallel double-stick tapes. The chamber was incubated with 100 nM anti-myo1b antibody (Almeida et al., 2011) in G buffer (5 mM Tris-HCl, pH 7.8, 0.1 mM CaCl_2_, 0.2 mM ATP, 1 mM DTT and 0.01% NaN_3_) for 10 min at room temperature. The chamber was rinsed three times with buffer G 0.1 % BSA and incubated 5 min at room temperature. Then the chamber was incubated with 300 nM Alexa488-labeled Myo1b in Fluo F buffer (5 mM Tris-HCl, pH 7.8, 100 mM KCl, 1 mM MgCl_2_, 0.2 mM EGTA, 0.2 mM or 2 mM ATP, 10 mM DTT, 1 mM DABCO, 0.01% NaN_3_) for 10 min at room temperature. Actin gliding assays were performed in Fluo F buffer, containing ATP maintained constant at 2 mM with a regenerating mix (see below), supplemented with 0.3% methylcellulose (Sigma) and with G-actin (1μM, 10% Alexa594-labelled), VCA and Arp2/3 complex or stabilized branched F-actin (stabilized with phalloidin-Alexa594) prepared as below. Branched F-actin was polymerized in presence of 1μM G-Actin, 100 nM VCA and 50 nM of Arp2/3 complex. In the case of stabilized filaments, after 10 min samples were diluted 20-fold to 50 nM and supplemented with phalloidin. The dynamic of stabilized branched filaments and polymerizing and branching actin filaments was monitored by TIRF microscopy (Eclipse Ti inverted microscope, 100X TIRF objectives, Quantem 512SC camera). The experiments were controlled using the Metamorph software. To maintaining a constant concentration of ATP in this assay an ATP regenerating mix, including 2 mM ATP, 2 mM MgCl_2_, 10 mM creatine phosphate and 3.5 U/mL creatine phosphokinase, which constantly re-phosphorylates ADP into ATP to maintain a constant concentration of free ATP, was added. Actin polymerization was measured as described by (Pernier et al., 2019) and expressed in actin subunit per second (su/s).

The sliding velocities of actin filaments were analyzed by using Kymo Tool Box plugin of Image J software (https://github.com/fabricecordelieres/IJ_KymoToolBox). The accuracy on the displacement of the filaments is of the order of the pixel size (160 nm). The number of branches and the length of the filaments were measured by using Analyze Skeleton plugin in Image J (https://imagej.net/AnalyzeSkeleton) on transformed movies obtained by running an automatic macro freely accessible on: http://xfer.curie.fr/get/SItjDQoyu4k/automatization_skeleton.ijm).

Statistical analysis was performed using Student t-test in Microsoft Excel.

## STORM

Stabilized branched F-actin in the absence of myosin or after 10 min of sliding on Myo1b was visualized by incubation with phalloidin-Alexa647. Samples were mounted in dSTORM buffer (Abbelight, Paris, France). Then, 3D nanoscopy images were taken using a SAFe360 module (Abbelight) coupled to the camera port of an inverted bright-field Olympus IX71 microscope, equipped with a 100×oil-immersion objective with a high numerical aperture (NA; 1.49). This quad-view system (dual-cam sCMOS cameras, Orcaflash v4, Hamamatsu) provided 3D nanoscopy information with high isotropic localization precision (15×15×15 nm, over an axial range of~1 mm). Axial information was obtained by super-critical angle fluorescence (SAF) and the point spread function (PSF) deformation with a strong astigmatism (DAISY) (Cabriel et al., 2018). Twenty-thousand frames at 50 ms were acquired to collect the single-molecule detections and reconstruct a nanoscopy image. Resulting coordinate tables and images were processed and analyzed using SAFe NEO software (Abbelight).

### Electron microscopy

4 μL of Myo1b or Myo2 samples was incubated on glow discharged and carbon coated electron microscopy grids (CF-300, EMS). After rinsing with the actin buffer, stabilized branched F-actin polymerized with 4 μM G-Actin, 100 nM VCA and 50 nM of Arp2/3 complex was incubated on the grid with 2 mM ATP. The sample was thereafter stained with 4 μL uranyl formate 2%. Images were collected with a Tecnai Spirit transmission electron microscope equipped with a LaB6 emission filament operated at an acceleration voltage of 80 kVolts (Thermofischer, FEI, Eindhoven, The Netherlands). The microscope is equipped with a Quemesa (Olympus) camera for imaging.

## Notes

### Competing Interest Statement

The authors have declared no competing interest.

